# Age-driven Dysregulation of murine Dendritic Cells is controlled by cell-intrinsic and extrinsic effects

**DOI:** 10.64898/2026.05.21.726941

**Authors:** Philipp Blöcher, Sunrito Mitra, Louise Guet, Emre Kilic, Zhilin Zou, Maximilian Sprang, Johannes U Mayer

## Abstract

Aging is associated with chronic, low-grade inflammation and progressive immune dysfunction. However, the current understanding of age-associated changes in dendritic cells across tissues is scarce. Studies exploring ageing-associated changes in dendritic cells (DCs) have reported either a general decline in the overall DC compartment or subset-specific alterations affecting cDC1, cDC2, and pDC populations across spleen, lung and liver, underscoring the considerable inconsistencies across tissues and studies. To underpin whether age-associated changes are extrinsic or intrinsic we investigated DCs across bone marrow, six peripheral tissues and in *in vitro* bone marrow derived DC cultures to examine the effects of aging on DC-poiesis, tissue distribution, and cellular states related to DC functionality and activation.

We discovered that aging selectively alters DC development in the bone marrow by reducing cDC progenitor populations while preserving pDC-poiesis. In peripheral tissues, however, age-associated changes in DC homeostasis were strongly tissue-dependent. The most significant shifts in cDC1 and cDC2 frequencies occurred in barrier tissues, such as the lung and small intestine. In contrast, the spleen and liver exhibited more limited or variable changes. These quantitative alterations were accompanied by tissue-specific changes in phenotypic and activation-associated markers, including CD24, CD103, CD11b, MHCII, and CD86. Single-cell transcriptomic analyses of senescent *p21*-expressing DC across tissues and subsets indicated localized inflammatory states that aligned with local macrophage populations, pointing toward cell-extrinsic niches that contribute to local age-associated dysfunction.

Notably, aged bone marrow retained the capacity to efficiently generate DCs in *in vitro* Flt3L cultures, and antigen-presenting function of BMDC to CD4 and CD8 T cells was maintained, pointing towards preserved cell-intrinsic functions, albeit subset-specific differences in activation and inhibitory receptor expression in response to different pattern-recognition receptor agonists.

Collectively, our findings indicate that aging does not superimpose a uniform alteration module to the DC compartment across tissues, but instead promotes selective alterations in DC ontogeny and tissue-specific remodeling of DC phenotypes and cellular states.

## Introduction

Aging is a multifactorial process characterized by a chronic low-grade inflammation (“inflammaging”) which has been associated with an increased risk for mortality and morbidity in the aged people, yet the cellular process remains incompletely understood^1,2^. From an immunological perspective, aging has been associated with a relative decline in lymphocytes and reciprocal increase in myeloid cells. Recent studies have linked this decline to impaired vaccination responses, associated a functional impairment of antigen-presenting cells within the myeloid compartment^3^. As major antigen-presenting cells, dendritic cells (DCs), which can be subdivided into plasmacytoid DCs (pDCs), conventional type 1 DCs (cDC1s), and conventional type 2 DCs (cDC2s), play central roles in coordinating innate and adaptive immune responses by their production of type I interferons^4^, antigen cross-presentation to cytotoxic CD8⁺ T cells^5^ and priming of naïve CD4⁺ T cells^6^.

Previous studies have elucidated that tissue-resident DC populations are continuously replenished from circulating precursors^7^. Thus, age-associated alterations in hematopoietic stem and progenitor cells (HSPCs) and DC progenitors may ultimately reshape the composition of local DC populations in a tissue-dependent manner. Previous studies have established that aging leads to an increased number of HSPC with a reduced differentiation potential, skewed differentiation toward the myeloid lineage and altered bone marrow (BM) homing and mobility^8,9^. These effects are shaped by the local BM niche, as adoptive transfer of old HSPC into a young irradiated recipient restores the transcriptional profile and capacity to generate myeloid cells, including cDCs, while leaving the aged HSPC intrinsic methylation profile largely unchanged^9,10^.

However, current reports on age-associated changes in DC distribution remain inconsistent between tissues and studies. For example, analyses of the spleen have described either a broad reduction in pDCs, cDC1s, and cDC2^11,12^ or, conversely, a significant increase in the cDC2 population^10,13,14^. Similarly, studies in the liver have reported divergent findings, ranging from increased pDC and cDC1 populations^15^ to a generalized reduction in pDC, cDC1 and cDC2^11^. Moreover barrier tissues display unique patterns^16–19^such as lung, where aging has been associated with reduced pDC and cDC1 frequencies, while cDC2 populations appear largely unchanged^11^. In the testis however, the cDC2 population remains stable whereas cDC1 population expands, while in the small intestine cDC1 frequencies remained unchanged^11^. Together, these findings suggest that aging-associated alterations in DC homeostasis are highly tissue-dependent and may be influenced not only by changes in precursor output but also by local microenvironmental cues. However, changes in DC abundance alone provides limited insight into their functional state within aged tissues.

Indeed, several studies have reported that aged DCs exhibit altered activation and maturation phenotypes in a tissue-specific manner, usually measured using markers such as CD86 and MHCII^20^. In the spleen, aged cDCs have been reported to exhibit increased CD86 expression under steady-state conditions^10^, while multiple other studies have observed no significant changes in MHCII and CD86 expression for cDCs and pDC^13,14,21^. Similarly, in the lung, aging was associated with increased MHCII expression in pDCs and cDC1s^15^ in some reports, but unchanged in others^10^. In addition aged cDC1s, in the liver, showed elevated MHCII, while CD86 and MHCII remained unchanged for the other DC populations^15^. Beyond alterations in classical activation markers, aging has also been associated with changes in innate signaling pathways in DCs, including reduced TLR7 expression and basal NF-κB activation in the spleen as well as increased lipid content in hepatic aged DCs^15,21^. Together, these findings indicate that aging does not induce a uniform DC activation state but rather promotes tissue-dependent inflammatory remodeling of DC function.

A major driver of tissue inflammaging is linked to the accumulation of senescent cells, a state of stable terminal cell-cycle arrest induced by diverse stressors. These cells contribute to inflammation via their senescent associated secretory phenotype (SASP) containing a mix of Interleukins, cytokines, chemokines, growth factors and extracellular matrix proteases^22^. SASP programs have been extensively characterized in stromal and parenchymal cell populations^23–25^ and also linked to tissue resident macrophages^26,27^. However, it remains unclear whether DCs also undergo senescence or express SASP associated genes. Recent transcriptomic atlas studies have revealed that aging induces both shared and cell type-specific transcriptional alterations, with the trajectory of aging being influenced by both cellular identity and tissue context^28^.

To reduce experimental complexity in the context of aging and test if cell-intrinsic age-associated DC alterations persist in the absence of the extrinsic tissue environment, bone marrow-derived dendritic cells (BMDC) have been used. GM-CSF/IL-4-driven BMDC cultures generated from aged mice yielded reduced numbers of differentiated DCs with increased basal expression of CD86, and MHC II under unstimulated conditions^29,30^. CCR7 expression was unchanged, but impaired migration toward CCL21 was observed *in vitro*^31^. Following LPS stimulation, however, reports are inconsistent, with some studies describing enhanced activation marker in aged BMDCs, while others observed reduced induction of MHC II and CD86 compared with young controls^29,30^. Furthermore, when GM-CSF IL-4 BMDCs were pulsed with OVA peptide_(257–264)_, no significant differences were observed between young and old in CD86 expression, nor in the uptake and surface presentation of OVA peptide–MHC I complexes. However reduced CD209 expression and a diminished capacity to induce T-cell proliferation was observed *in vitro*^31^, yet it remains unclear how Flt3L (Fms-like tyrosine kinase 3 ligand) differentiated BMDC, which more resemble cDC than moDC^32,33^, behave in these contexts.

To more closely define the age-associated changes of conventional cDCs we deployed detailed flow cytometric profiling of DC and DC precursors across the bone marrow and six peripheral tissues and found that DC-poiesis, as well as tissue distribution and senescence-associated profiles, were affected in a tissue and subset dependent manner. In vitro, DC activation was however only marginally changed across a wide array of agonists with primary DC functions linked to T cell priming remaining unchanged, indicating that changes within the aged tissue environment imprint local DC phenotypes, rather than cell-intrinsic defects.

## Results

### Aging induces a decline of DC-poesis in murine bone marrow

To examine the impact of aging on DC-poiesis, we first curated a unified ontogenetic framework from the existing literature to systematically resolve the stepwise differentiation of HSPCs into DCs. As the earliest progenitors within the DC developmental hierarchy, HSPCs reside in the Lin^-^, Sca-1^+^, cKit^+^ (LSK) compartment and can be further subdivided into distinct hematopoietic stem and multipotent progenitor populations according to Sommerkamp *et al.* These include HSC (LSK CD34^-^,CD48^-^,CD150^+^,CD135^-^), MPP1 (LSK CD34^+^,CD48^-^,CD150^+^,CD135^-^), MPP2 (LSK CD34^+^,CD48^+^,CD150^+^,CD135^-^), MPP3 (LSK CD34^+^,CD48^+^,CD150^-^,CD135^-^), MPP4 (LSK CD34^+^,CD48^+^,CD150^-^,CD135^+^), MPP5 (LSK CD34^+^,CD48^-^,CD150^-^,CD135^-^) and MPP6 (LSK CD34^-^,CD48^-^,CD150^-^,CD135^+^)^34^. Our flow cytometric analysis from bone marrow of young (8 weeks of age) and old (72–73 weeks of age) male C57BL/6J mice revealed a significant age-associated increase in the frequencies of HSCs and MPP1 cells (Fig. 1b), populations previously reported to reside predominantly in the G0/G1 phase and represent the most upstream compartments of hematopoiesis. In parallel, aging was associated with a significant reduction in both the myeloid-biased MPP3 and lymphoid-biased MPP4 populations (Fig. 1b), whereas the frequencies of MPP2, MPP5 and MPP6 remained unchanged.

**Fig 1.**
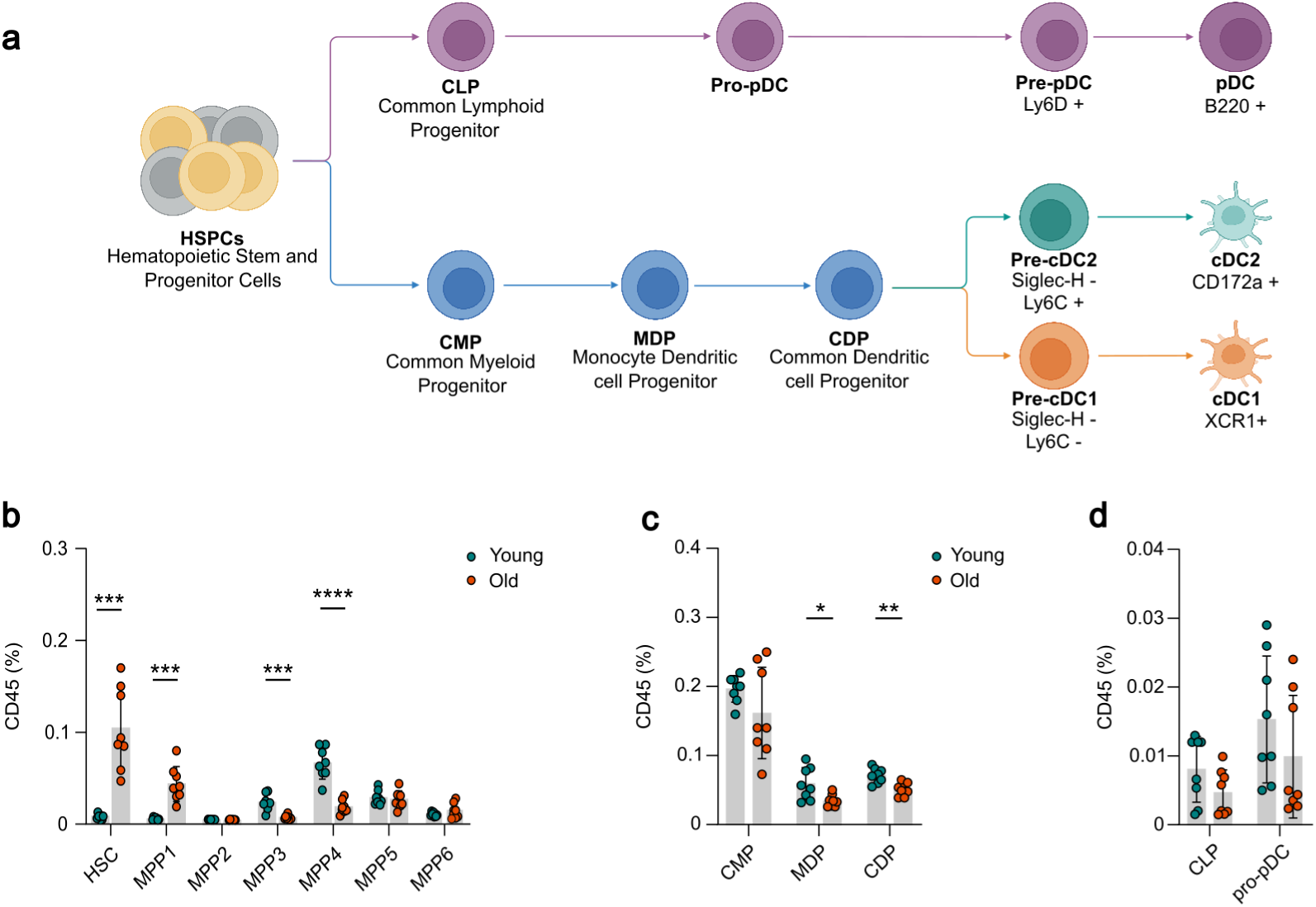
Aging associated alterations of cDC-poiesis in bone marrow. **a**, Schematic overview of DC-poiesis consisting of myeloid-derived cDC-poiesis (blue) with cDC1- (orange) and cDC2-committed (green) precursors and lymphoid–derived pDC-poiesis (purple). **b**, Age-dependent relative abundance of bone marrow LSK cell populations, particularly HSC, MPP1, MPP2, MPP3, MPP4, MPP5 and MPP6. **c**, Frequencies of multipotent progenitors of cDC-poiesis in young and old mice. **d**, Proportions of pDC-poietic non-committed progenitors. **b-d**, Bar graphs show mean ± s.d. for n=8 mice pooled from two independent experiments. All experiments used young (8 weeks) and old (72–73 weeks) C57BL/6 mice. Each symbol represents one mouse. *P* values were determined using unpaired two-tailed t tests with Holm-Sidak correction. * *P* <0.05, ** *P* <0.01, *** *P* <0.001, **** *P* <0.0001; only significant comparisons are indicated.

To further investigate DC-poiesis downstream from the HSPCs we analyzed the myeloid progenitor compartment and specifically DC precursors (Fig. 1a). cDCs originate from the CD135^+^ common myeloid progenitor (CMP) (Lin^-^, Sca-1^-^, cKit^+^, CD34^+^, CD16/32^lo^, CD115^-^, CD135^+^) which gives rise to the monocyte dendritic cell progenitor (MDP) (Lin^-^, Sca-1^-^, cKit^+^, CD34^+^, CD16/32^lo^, CD115^+^, CD135^+^)^35^. MDPs subsequently differentiate through CD115+ common dendritic cell progenitors (CDPs) (Lin^-^, CD11c^-^, MHCII^-^, CD135^+^, CD117^int^, CD115^+^) into the pre-cDC (Lin^-^, CD11c^+^, MHCII^-^, CD172a^-^,CD135^+^, Ly6D^-^), which divide into committed pre-cDC1 (Siglec-H^-^Ly6C^-^) and pre-cDC2 (Siglec-H^-^Ly6C^+^) ^35,36^, and finally into cDCs (Lin^-^, CD11c^+^MHC II^+^CD26^+^). They can be further subdivided into XCR1^+^CD24^+^cDC1 and CD11b^+^CD172a^+^ cDC2^37^. In contrast, pDCs (Lin^-^, B220^+^) appear to be primarily lymphoid-derived, originating from the Ly6D+ subpopulation of the common lymphoid progenitor (CLP) (Lin^-^, CD127^+^, cKit^lo^, Sca-1^lo^, CD135^+^, CD34^+^, Ly6D^+^) via a CD115^-^Ly6D^+^ CDP-like pro-pDC (Lin^-^, CD11c^-^, MHCII^-^, CD135^+^, CD117^int^, CD115^-^, Ly6D^+^) and Ly6D^+^ pre-pDCs (Lin^-^, CD11c^+^, MHCII^-^, CD172a^-^,CD135^+^, Ly6D^+^)^38^.

Within DC precursors we observed a significant reduction in CD115^+^ CDP and MDP cells, which was also observed as a trend in CD135^+^ CMP cells (Fig. 1c). Within the lymphoid branch, CD45% frequencies of both Ly6D^+^ CLP and Ly6D^+^ pro-pDC pDCs were comparable in the bone marrow of young and old mice (Fig. 1d), indicating that myeloid derived DC precursor frequencies are stronger impacted in aged mice than lymphoid derived pDC precursors.

### cDC1 in aged tissues show subtle changes in distribution and transcriptional programs mainly affecting small intestine and testis

To investigate whether ageing remodels the cDC1 compartment across tissues, we first quantified cDC1-committed pre-DCs in the bone marrow, defined as Ly6D⁻Siglec-H⁻Ly6C⁻CD11c⁺MHCII⁻CD135⁺CD172a⁻ (Fig. 1a). The relative frequency of this population was comparable between young and aged mice, reflecting no marked impairment in the generation of cDC1-committed precursors in the bone marrow (Fig. 2a).

**Fig 2.**
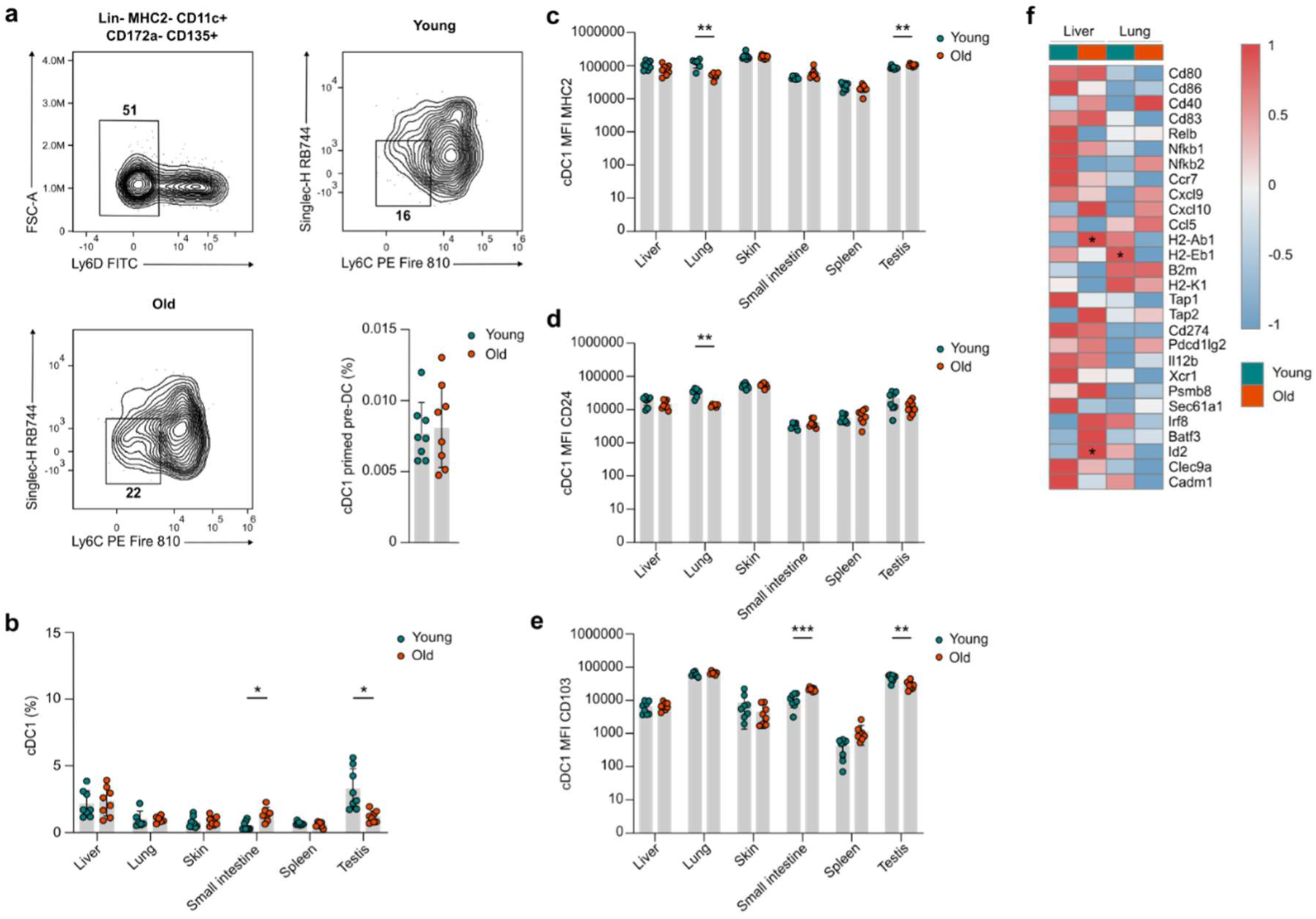
Aging affects the cDC1 compartment in certain murine tissues. **a**, Flow cytometric analysis of SiglecH-Ly6C- pre-cDC1 in bone marrow. Representative flow cytometry plots and frequencies for young and old mice are shown. **b**, Frequencies of XCR1+ cDC1 cells in peripheral tissues including liver, lung, skin, small intestine, spleen, and testis of young and old mice. **c**, Median fluorescence intensity (MFI) of MHCII within cDC1 population across indicated tissues. **d**,**e** MFI of CD24 and CD103 within cDC1 population across indicated tissues. **f**, Heatmap of differentially expressed genes between young and old DC1 in liver and lung. Z-scores for each gene were calculated using R. **a-e**, Bar graphs show mean ± S.D. for n = 8 mice (**a**) or n = 6-8 mice (**b-e**) pooled from two independent experiments. All experiments used young (8 weeks) and old (72–73 weeks) C57BL/6 mice. Each symbol represents one mouse. Each symbol represents one mouse. *P* values were determined using unpaired two-tailed t tests with Holm-Sidak correction (**a-e**) or Wilcoxon rank-sum test (**f**). For panels **a**-**e**: \**P* <0.05, ** *P* <0.01, *** *P* <0.001; only significant comparisons are indicated.

We next examined mature cDC1s across peripheral tissues. In aged mice we observed a selective reduction in cDC1 abundance in the testis, while a significant increase was observed in the small intestine, with comparable frequencies in the spleen, liver, lung and skin (Fig. 2b).

With the extensive and established phenotypic heterogeneity of tissue-resident cDCs already published^39^, we next assessed whether ageing altered cDC1 identity or activation-associated marker expression. Analysis of CD24 and CD103 revealed marked tissue-specific changes in aged mice. CD24 expression was selectively reduced on lung cDC1s from aged mice (Fig. 2d). In the small intestine, DC1 from aged mice showed increased CD103 expression levels (Fig. 2e), whereas testicular cDC1s showed reduced CD103 expression (Fig. 2e), showing parallel patterns between CD103 expression and cDC1 frequency. DC1 from aged mice further showed a significant increase in MHCII expression in the testis and lung (Fig. 2c). On a transcriptomic level we re-analyzed data from a publicly available scRNA-seq dataset^40^ from the liver and lung of young (6-8 weeks of age) and aged (72 weeks of age) mice, defined DC1 within the dendritic cell cluster through their characteristic expression of *Irf8*, *Id2* and *Plbd1* and assessed transcripts related to DC1 activation and identity (Fig. 2f). Lung DC1 from aged mice showed a significant decrease in *H2-Eb1*, consistent with our flow cytometry data. On the transcriptomic levels of *H2-Ab1* were significantly increased in liver DC1 from aged mice together with and *Id2*, yet MHC2 surface marker expression was unchanged in liver DC1 of young and old mice, based on our flow cytometric analysis (Fig. 2c). While several non-significant changes were observed, normalizing expression levels between DC1 from different tissues, also revealed striking differences in canonical DC1 genes between the liver and lung in general, with the lung trending towards lower expression levels overall.

### cDC2 frequency is reduced in aged lungs and testes and shows increased activation in specific tissues

We similarly quantified cDC2-committed pre-DCs in the bone marrow, defined as Ly6D⁻Siglec-H⁻Ly6C⁺CD11c⁺MHCII⁻CD135⁺CD172a⁻ (Fig. 1a). In contrast to cDC1-committed precursors, the frequency of cDC2-committed pre-DCs was significantly reduced in aged mice, suggesting a potential decline in cDC2-poiesis with ageing (Fig. 3a).

**Fig 3.**
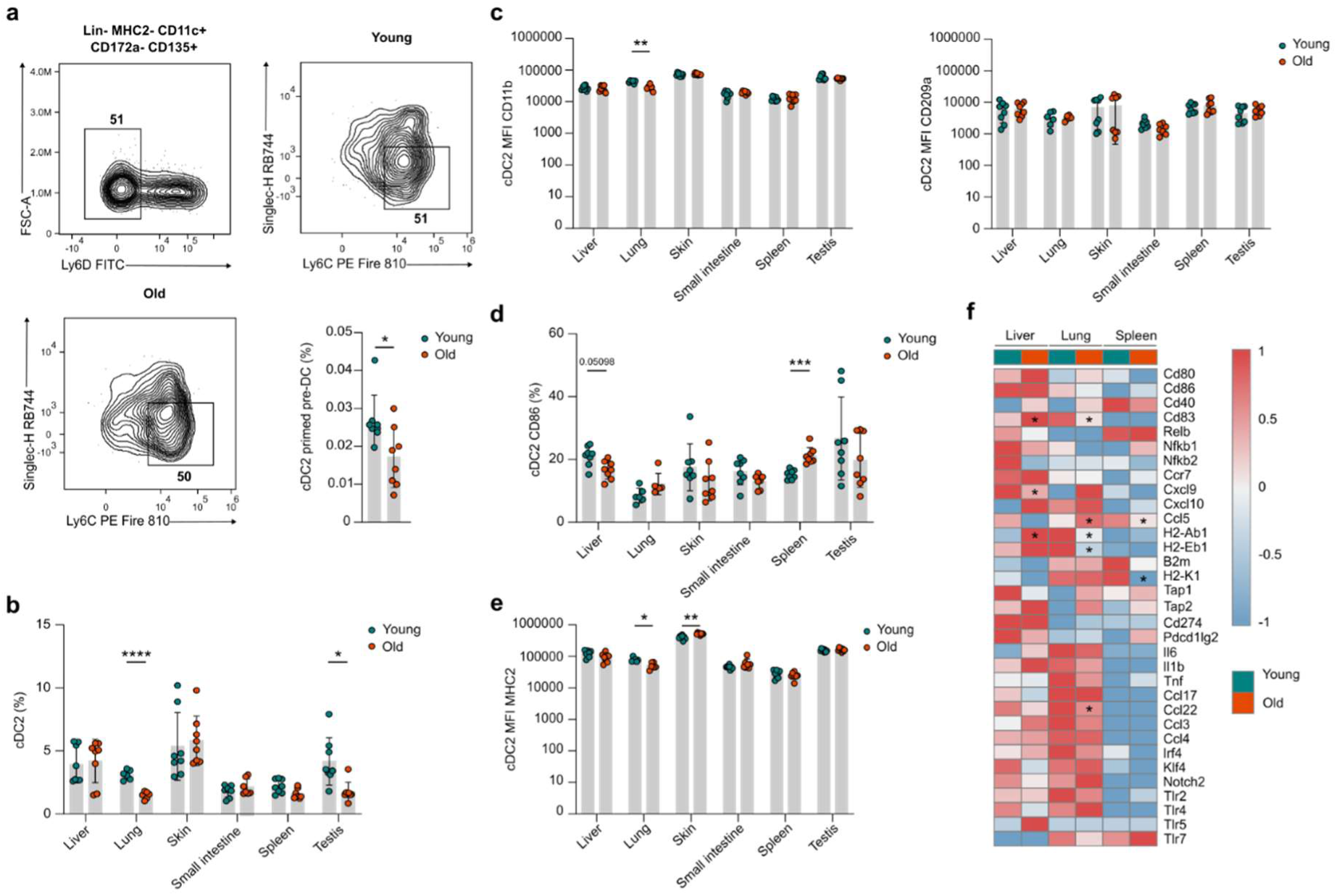
cDC2s exhibit heterogeneous aging patterns in different murine tissues. **a**, Flow cytometric analysis of SiglecH-Ly6C+ pre-cDC2 in bone marrow. Representative flow cytometry plots and frequencies for young and old mice are shown. **b**, Frequencies of CD172a+ cDC2 cells in peripheral tissues including liver, lung, skin, small intestine, spleen, and testis of young and old mice. **c**, Median fluorescence intensity (MFI) of CD11b and CD209a within the cDC2 population across indicated tissues. **d**,**e** Frequencies of CD86+ cDC2 cells and MFI of MHCII within cDC2 population across indicated tissues. **f**, Heatmap of differentially expressed genes between young and old clusters in liver, lung and spleen. Z-scores for each gene were calculated using R. **a-e**, Bar graphs show mean ± S.D. for n = 8 mice (**a**) or n = 6-8 mice (**b-e**) pooled from two independent experiments. All experiments used young (8 weeks) and old (72–73 weeks) C57BL/6 mice. Each symbol represents one mouse. Each symbol represents one mouse. *P* values were determined using unpaired two-tailed t tests with Holm-Sidak correction (**a-e**) or Wilcoxon rank-sum test (**f**). For panels **a**-**e**: \**P* <0.05, ** *P* <0.01, *** *P* <0.001; only significant comparisons are indicated.

Mature cDC2 populations were then assessed across peripheral tissues. cDC2 were significantly reduced in lung and in testis, while remaining unchanged in liver, skin, small intestine, and spleen (Fig. 3b). To evaluate if aging changes cDC2 tissue phenotypic heterogeneity, we looked into the tissue specific markers such as CD11b and CD209a^31,41^. We observed a significant reduction of CD11b expression levels in the lung, but not in other tissues, with CD209 expression levels also being unchanged (Fig. 3c). Instead, we observed a modest increase in the proportion of CD86⁺ activated cDC2 in spleen, and a decreasing trend in liver, small intestine and testis (Fig. 3d). Overall expression levels of MHCII were reduced in lung cDC2, while increased in skin cDC2 (Fig. 3e), indicating that frequencies and MFI expression levels do not necessary correlate with each other.

On a transcriptomic level, re-analysis of DC2 from a publicly available scRNA-seq dataset^40^ from young (6-8 weeks of age) and aged (72 weeks of age) mice, showed that in the liver, aged cDC2 express increased levels of *H2-Ab1* and *Cd83* alongside decreased levels of *Cxcl9*, suggesting increased activation in this dataset (Fig. 3f). In line with our flow cytometric analysis, lung cDC2 showed reduced activation with age, including decreased expression of *H2-Eb1*, *Cd83*, and *H2-Ab1* (Fig. 3f). Overall, when comparing DC2 from different tissues, splenic cDC2 showed the lowest expression of canonical cDC2 genes with altered cDC2 activation being observed in a tissue-specific manner.

### Age-related reduction of pre-pDCs in the bone marrow correlates with reduced pDC frequencies in liver

In line with previous publications, aged mice exhibited a significant reduction in pre-pDC frequency, defined as the Lin^-^, CD11c^+^, MHCII^-^, CD172a^-^,CD135^+^, Ly6D^+^, suggesting an age-associated impairment in pDC commitment and/or maintenance within the bone marrow (Fig. 4a).

**Fig 4.**
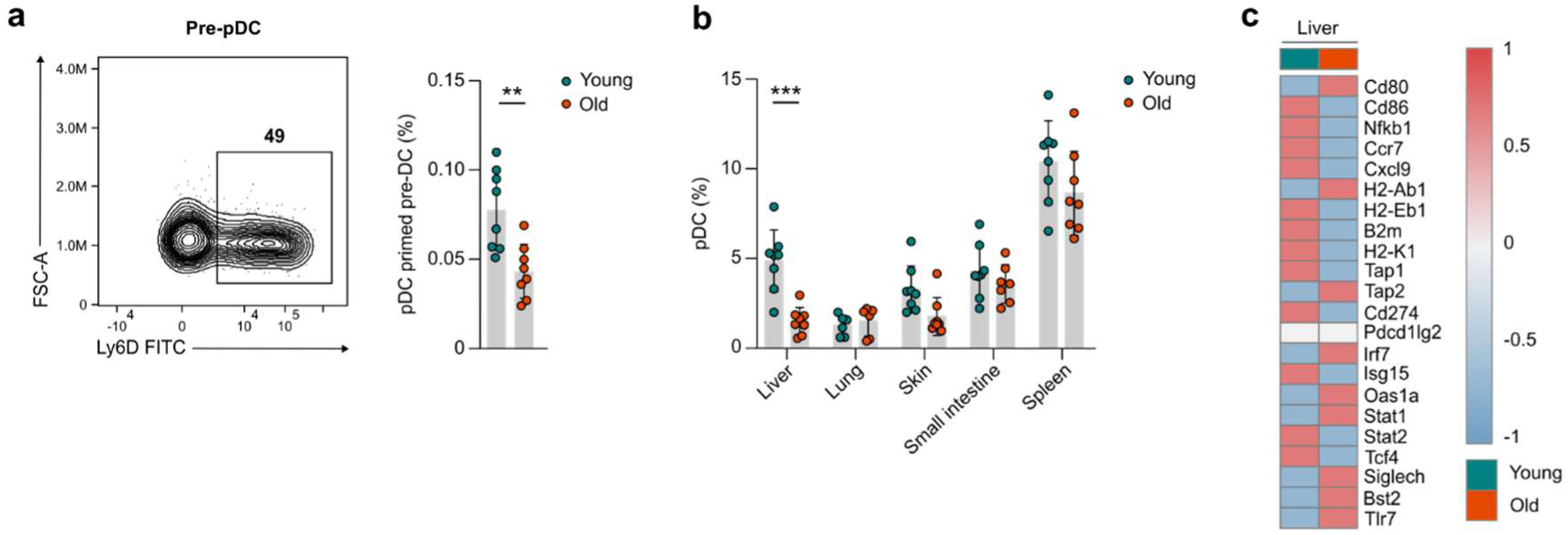
Age-related reduction of pre-pDCs in the bone marrow correlates with reduced pDC frequencies in liver. **a**, Flow cytometric analysis of Ly6D+ pre-pDC in bone marrow. Representative flow cytometry plot and frequencies for young and old mice are shown. **b**, Frequencies of B220+pDC in peripheral tissues including liver, lung, skin, small intestine, and spleen of young and old mice. **c**, Heatmap of differentially expressed genes between young and old clusters in liver. Z-scores for each gene were calculated using R. **a-b**, Bar graphs show mean ± S.D. for n = 8 mice (**a**) or n = 6-8 mice (**b-e**) pooled from two independent experiments. All experiments used young (8 weeks) and old (72–73 weeks) C57BL/6 mice. Each symbol represents one mouse. Each symbol represents one mouse. *P* values were determined using unpaired two-tailed t tests with Holm-Sidak correction (**a,b**) or Wilcoxon rank-sum test (**f**). For panels **a,b**: ** *P* <0.01, *** *P* <0.001; only significant comparisons are indicated.

Within peripheral tissues, pDC frequencies were selectively decreased in the liver of aged mice, unchanged in other tissues and not detected in the epididymis or testis, which aligns with previous studies^42^ (Fig. 4b). pDC-associated gene expression signatures were also not significantly altered in the liver of young (6-8 weeks of age) and aged (72 weeks of age) mice in a publicly available scRNA-seq dataset^40^, indicating that the core transcriptional identity of hepatic pDCs is largely preserved during ageing despite reduced cellular abundance (Fig. 4c). Together, these data suggest that ageing diminishes pDC output at the precursor stage and selectively reduces hepatic pDC frequency, with limited effect to the transcriptional programme of the remaining pDC population.

### Altered TLR agonist responsiveness and PD-L1 and PDL-2 upregulation are features of BMDC cultured from aged bone marrow

To detect cell-intrinsic alterations of DC functionality, we cultured BMDC from the BM of old and young mice under Flt3L conditions. Assessment of the *in vitro* differentiation potential of HSPCs from young and old BM showed no significant age-associated differences in the output of pDCs, cDC1s or cDC2s (Fig. 5a). Under unstimulated conditions, BMDC subsets displayed distinct basal expression patterns of CD86 and PD-L1, which cDC1s showing high levels of CD86 and cDC2s showing a distinct PD-L1 expression pattern (Fig. 5b). However, these profiles were not significantly altered in BMDCs generated from aged mice compared with young controls (Fig. 5b). Following LPS stimulation, we observed marked upregulation in activation-marker expression patterns across DC subsets. cDC1s expressed elevated levels of CCR7, CD40, CD86, PD-L1 and PD-L2, whereas cDC2s modestly expressed CCR7 and PD-L1, but showed high expression of CD40 and CD86, with low expression of PD-L2, with pDCs responding little to LPS stimulation (Fig. 5b).

**Fig 5.**
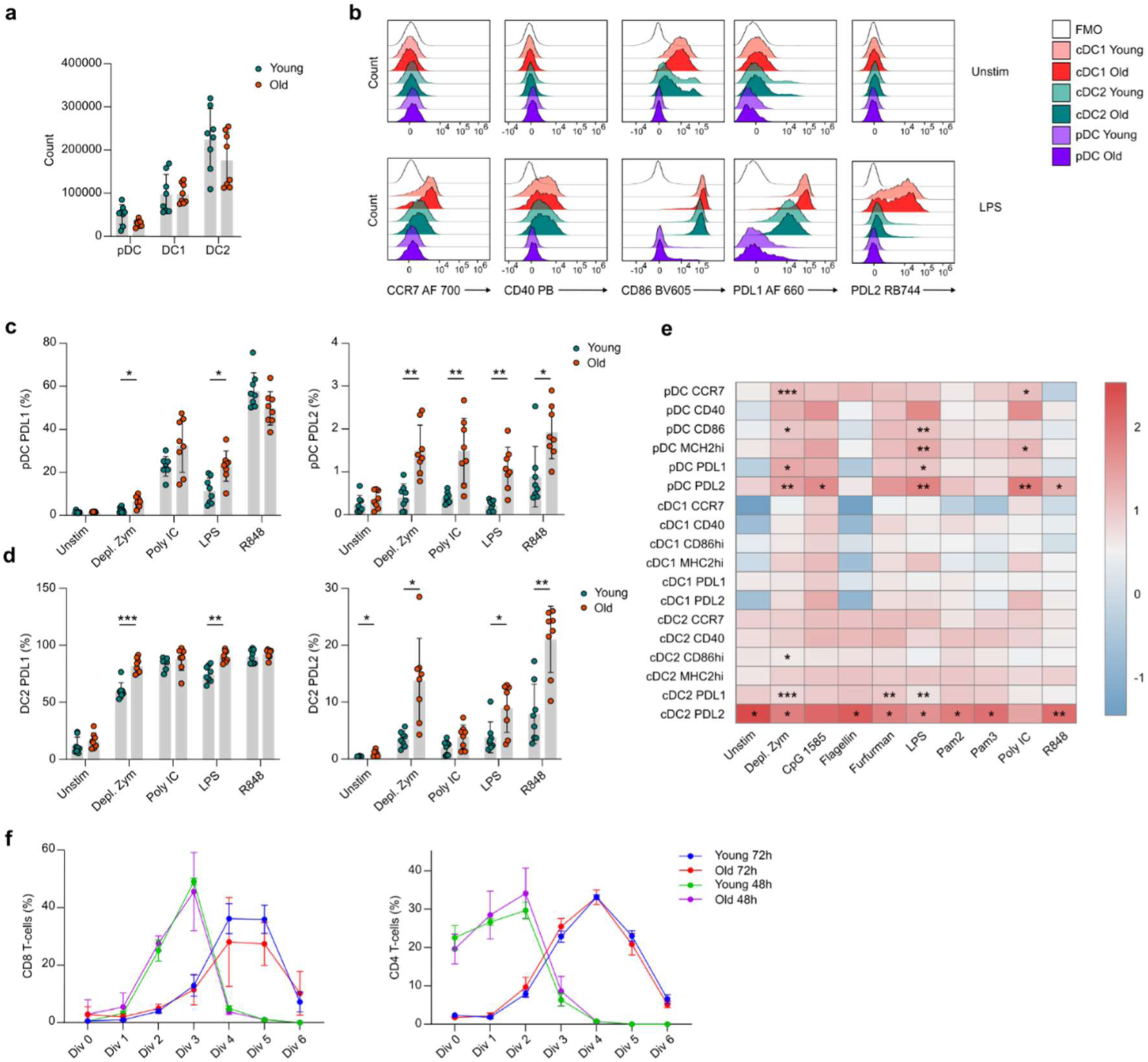
Altered TLR agonist responsiveness and PD-L1 and PDL-2 upregulation are features of BMDC cultured from aged bone marrow. **a**, Counts of DC subsets in BMDC per 106 input cells from young vs old bone marrow. **b**, Representative Histograms of indicated activation and inhibition marker expression patterns from young and old BMDC subsets with or without LPS stimulation (5 ng ml-1 for 18 h). **c**,**d** Selected bar graphs showing expression of PD-L1 and PD-L2 on pDC and cDC2 in response to indicated TLR agonists. **e**, Heatmap of differentially expressed activation and inhibitory markers (Log2 fold change old to young) measured by flow cytometry in young and old BMDC subsets in response to indicated TLR agonists. **f**, Line graphs showing percentage of OT-1 and OT-2 cell divisions after 48 h and 72 h co-culture with BMDC pulsed with OVA (200 µg mL-1 for 18 h). Values represent the mean of two technical replicates, with n = 8 mice per group. Data are presented as mean ± S.D. **a**,**c**,**d**, Bar graphs show mean ± S.D. for n = 8 mice pooled from two independent experiments. All experiments used young (8 weeks) and old (72–73 weeks) C57BL/6 mice. Each symbol represents one mouse. *P* values were determined using unpaired two-tailed t tests with Holm-Sidak correction (**a,c-e**). For panels (**a,c-e**): \**P* <0.05, ** *P* <0.01, *** *P* <0.001; only significant comparisons are indicated.

Age-associated changes were particularly pronounced in cDC2s and pDCs, particularly involving CD86, PD-L1 and PD-L2 expression, while cDC1 showed no differences. Notably, PD-L2 expression was increased in aged cDC2s under unstimulated conditions and after stimulation with depleted zymosan, LPS or R848, which engage Dectin-1, TLR4 and TLR7, respectively (Fig. 5d). An increase in PD-L1 expression was also observed in aged cDC2s after depleted zymosan and LPS stimulation with CD86 expression also being higher in BMDC from aged mice after Dectin-1 stimulation with depleted zymosan (Fig. 5d).

Aged BMDC-derived pDCs also exhibited significant higher levels of CCR7, CD86, PD-L1 and PD-L2 compared to their young BM counterparts (Fig. 5c). Similar to aged cDC2s, pDCs showed increased PD-L1 expression following stimulation with depleted zymosan or LPS (Fig. 5c). In addition, PD-L2 expression was elevated in aged pDCs after stimulation with depleted zymosan, LPS, poly(I:C) and R848, corresponding to Dectin-1, TLR4, TLR3 and TLR7 activation, respectively (Fig. 5c). These changes in costimulatory marker profiles did however not affect the priming of CD4 or CD8 T cells when BMDCs from young or old BM were pulsed with OVA and subsequently co-cultured with MACS-purified OT-I or OT-II T cells for 48 h or 72 h, as no changes in the proliferation of CD4 and CD8 T cells was observed (Fig. 5f).

### Senescence associated *p21*-expressing DCs display independent transcriptional programs across tissues

In the context of inflammaging senescent cells have been shown to play an important role as they accumulate within tissue niches and lead to a proinflammatory milieu. *p21*, also known as *Cdkn1a*, has been established as a senescence marker in RNA-seq studies^22,23^, recently described to be associated with a senescent phenotype in liver macrophages. DCs and macrophages in the followed a similar p21-experssion trajectory over different ages in our curated dataset GSE153562^27,40^, which was observed across DC subsets (Fig. 6a,b).

**Fig 6.**
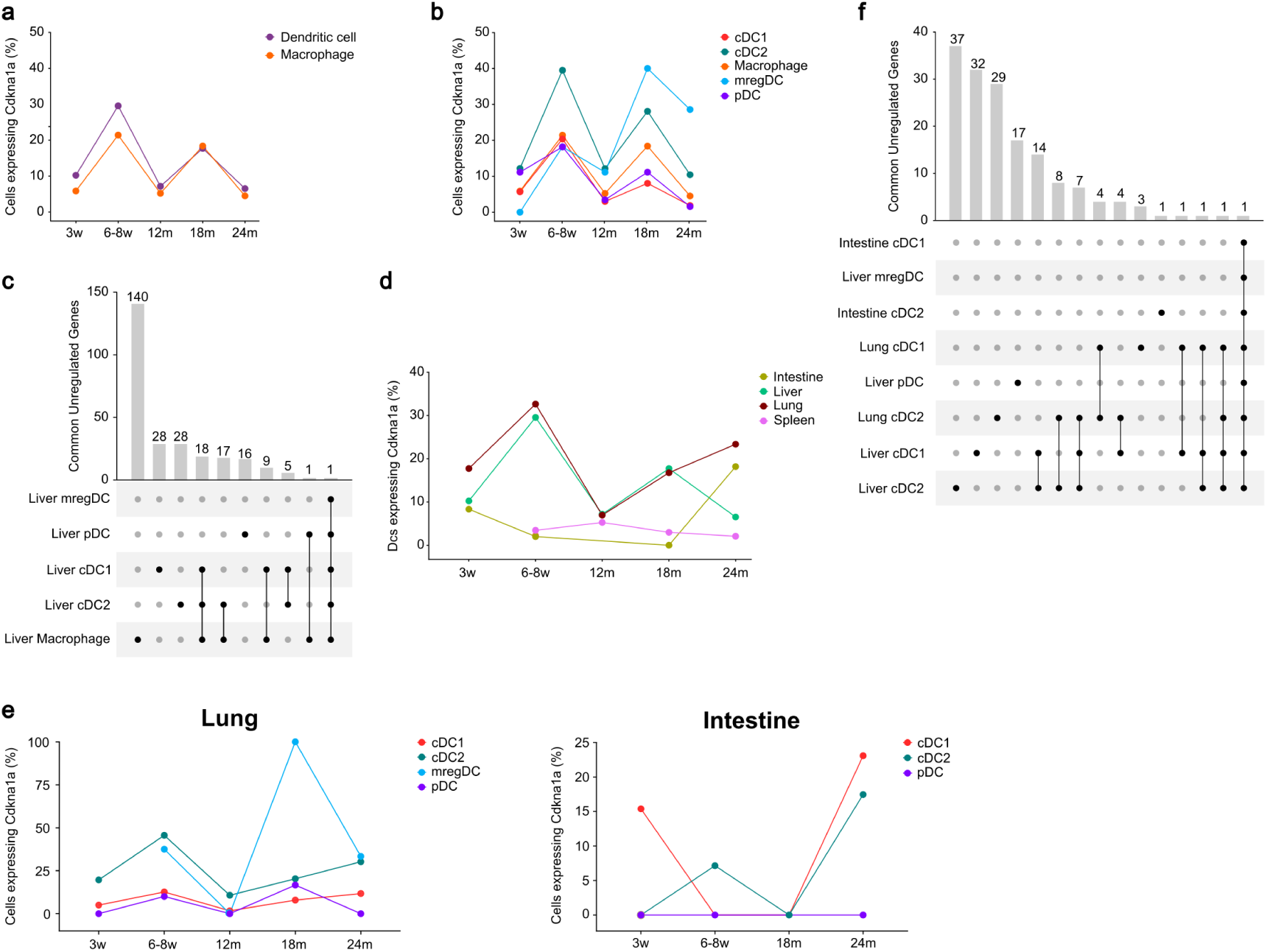
*p21*-expressing cDCs display distinct transcriptional signatures across subsets and tissues. **a**, Percentage of *p21*-expressing macrophages and dendritic cells in liver at different age time points in dataset GSE15356240. **b**, Percentage of *p21* expression across DC subsets (pDC, cDC1, cDC2, migDC) and macrophages in liver in dataset GSE15356240. **c**, Upset plot showing overlapping differentially expressed genes (DEGs) upregulated in *p21+* versus *p21-* macrophages and DC subsets in liver in dataset GSE15356240. **d**, Percentage of *p21*-expressing dendritic cells in liver, lung, spleen and intestine at different age time points in dataset GSE15356240. **e**, Percentage of *p21* expression across DC subsets (pDC, cDC1, cDC2, migDC) in lung and intestine in dataset GSE15356240. **f**. Upset plot showing overlapping DEGs of *p21+* versus *p21-* upregulated genes for DC subsets across liver, lung and intestine in dataset GSE15356240.

Liver macrophages expressed a high number of differentially expressed genes (DEG) between *p21*+ and *p21*- populations, while DC expressed much fewer DEGs, many of which were shared between different subsets and macrophages (Fig. 6c). While liver and lung showed comparable proportions of *p21* expressing DCs up to 18 months, intestine and spleen showed overall less *p21^+^* cells (Fig. 6d,e). Across tissues, DC subsets showed divergent *p21*-associated DEG patterns with only few overlapping genes (Fig. 6f). cDCs in lung and liver both shared *Pim1* upregulation in the cDC compartment, an inducer of *NF-κB*^43^, *Cd14*, a TLR4 co-receptor, as well as the proinflammatory cytokine *Il1b* (Fig. 6f), which is in contrast to the much larger shared overlap between senescent macrophages and non-immune cells across tissues^23,27^.

## Discussion

Here, we report that DC-poiesis shows subset specific alterations in the cDC precursor and subset pool. The frequencies of cDC1 and pDC precursors remain largely unchanged within the bone marrow of aged mice, while cDC2 population were reduced. In our hands, myeloid precursor output did not increase dramatically in aged bone marrow. Rather, lymphoid progenitor cell were reduced, corroborating published findings that reported a shift towards monopoiesis with aging^11,44^.

Within tissues, we observed changes in distribution and activation marker profiles across multiple cDC-subsets, which were particularly prominent in barrier tissues and the testis. In the immunoregulatory environment of the testis^45^, we observed a reduction in cDC1 numbers as well as a decreased CD103 expression with aging, DC types associated with Treg programs in other tissues^46^. The heterogeneity observed across testis, small intestine and lung, further indicates that DC aging cannot be explained by a uniform systemic program, but instead reflects organ-specific signature and reprograming.

Previous studies have reported age-associated alterations in DC subsets; however, findings have been inconsistent across tissues and subsets ^10,11^. These discrepancies potentially also reflect differences in DC subset definitions and gating strategies, particularly in earlier work relying on CD11b for cDC2 and CD8α/CD103 expression for cDC1. In contrast, our study employed state-of-the-art DC classification strategies with XCR1 defining cDC1 and Sirpa defining cDC2^47,48^. Recent studies have also defined an additional DC population within the Sirpa+ compartment^49^, however due to a lack of verified gating strategies in tissues, DC3 were not studied in this context.

*In vitro,* aged BMDC cultures exhibited largely preserved baseline DC output and steady-state phenotype, suggesting that core differentiation capacity remains intact during aging, unlike GM-CSF/IL-4 BMDC cultures, which depict an increased activation profile^29,30^. In addition, we observed that aged BM-pDCs and cDC2s showed a stronger activation and PD-L1 and PD-L2 expression profile, upon stimulation of TLR4 and Dectin-1, suggesting an increased T cell activation and inhibitory program^50,51^. While in *in vitro* cocultures OTI and OTII T cell proliferation was unchanged between BMDC cultured from young and old BM, tissue specific signals might still alter local DC priming capacity in aging.

Local senescent cell accumulation and senescence-associated secretory phenotype (SASP) are established drivers of inflammaging and observed in the non-hematopoietic compartment and macrophages^27^. Our analysis of *p21-*expressing cDC across tissues, showed that cDC express a much lower number of *p21*-associated DEGs compared to macrophages, but display tissue and subset-specific profiles that might influence highly context dependent local environments^28^.

Together, these findings support a model in which dendritic cell aging is not defined by a uniform loss of function or systemic decline in developmental capacity, but instead suggest context-dependent reprogramming. BM DC-poiesis is selectively altered, while core Flt3L-driven differentiation capacity remains largely intact. In addition, tissue environments might regulate DC phenotypes, leading to some of the observed changes in DC phenotypes, especially in barrier tissues. Within these tissues, DCs can acquire context-dependent activation states, either through cell-extrinsic or cell-intrinsic *p21*-associated inflammatory programs.

## Acknowledgements

The authors gratefully acknowledge the data storage facilities provided by the Institute for Quantitative and Computational Biosciences (IQCB) at Johannes Gutenberg University (JGU) Mainz. JUM acknowledges funding from the ReALity Initiative of the Johannes Gutenberg Universität Mainz and the Forschungsinitiative des Landes Rheinland-Pfalz, the Rise up! programme of the Boehringer Ingelheim Foundation (BIS), German Research Foundation (DFG) Research Unit FOR 5644, project no.515636567 and German Research Foundation (DFG) Research Training Group GRK 2573/2. PB acknowledges funding from the “ImmunYoAge” Else Kröner Promotionskolleg. The authors extend their gratitude to Tina Sarkar, Bingbing Chen, Jeevan Mutha, André Heine, Emma-Lou Bökenkamp and Sanjana Zende for their experimental support.

## Author contributions

P.B.: Conceptualization, Data curation, Formal analysis, Investigation, Methodology, Visualization, Writing – original draft; S.M.: Data curation, Formal analysis, Visualization, Writing – review & editing; L.G.: Data curation, Formal analysis, Visualization; Z.Z.: Software, Methodology, Writing – review & editing; E.K.: Software, Methodology, Writing – review & editing; M.S.: Software, Methodology, Writing – review & editing; J.U.M.: Conceptualization, Funding acquisition, Project administration, Resources, Supervision, Writing – review & editing.

## Materials and Methods

### Animals

All animals in this study were of C57BL/6J background and were either bred and housed at the Translational Animal Research Center (TARC) of the Johannes Gutenberg University of Mainz according to institutionally approved protocols, or purchased from Charles River Laboratories and acclimatized at the TARC for at least 2 weeks before experimental procedures. Animal experiments were conducted under the supervision of authorized investigators in accordance with European Union regulations for the care and use of experimental animals and all relevant ethical guidelines. Experiments used young (8 weeks of age) and old (72–73 weeks of age) male C57BL/6J mice. Male and female B6 OT-I × B6 Ly5.1 and B6 OT-II × B6 Ly5.1 mice (36-94 weeks of age) were kindly provided by Dr. Matthias Bros and Prof. Stephan Grabbe from the Department of Dermatology, Mainz and bred at the TARC.

### Tissue harvesting and single-cell suspension preparation

#### Liver perfusion and single-cell preparation

Similar to Ponti et al.^52^, murine livers were perfused in situ prior to excision with collagenase A (0.2 mg/mL; Roche / 10103586001) in IMDM (Sigma Aldrich / I3390) to reduce blood content. Tissue was mechanically dissociated and further digested in 2 mL RPMI (Sigma Aldrich / R8758) containtg collagenase A (0.4 mg/mL) and DNase I (0.01 mg/mL; Sigma Aldrich / 10104159001) at 37 °C for 15 min with intermittent mixing. After centrifugation at 400g, for 5min cells were washed in PBS. The cells were then filtered through a 100 µm strainer and treated with RBC lysis buffer (BioLegend / 420302) at 4°C. Samples were subsequently filtered through a 40 µm strainer to remove aggregates and processed for counting and flow cytometry.

#### Spleen single-cell preparation

Murine spleens were harvested, mechanically dissociated by mincing, and digested in 1 mL RPMI containing Liberase TL (0.1 mg/mL; Sigma Aldrich / 5401020001) and DNase I (0.05 mg/mL) for 30 min at 37 °C in a thermomixer. Cell suspensions were filtered through a 70 µm strainer, followed with a RBC lysis. Cells were washed, resuspended in FACS buffer, and used for counting and downstream flow cytometry analysis.

#### Lung single-cell preparation

Murine lungs were harvested and half lungs were mechanically dissociated by mincing, followed by enzymatic digestion in 1 mL RPMI containing Liberase TL (0.5 mg/mL) and DNase I (0.5 mg/mL) for 45 min at 37 °C in a thermomixer (600 rpm). Following filtration through a 70 µm strainer, red blood cells were lysed, and used for counting and downstream flow cytometry analysis.

#### Lymph node and single-cell preparation

Murine lymph nodes were harvested, mechanically dissociated using 18G needles and digested in 1 mL RPMI containing Liberase TM (100 µg/mL) and DNase I (100 µg/mL) for 30 min at 37 °C on a thermomixer. Cell suspensions were then filtered through a 70 µm strainer and processed for downstream analysis and cell counting.

### Small intestinal lamina propria single-cell preparation

Small intestinal lamina propria cells were isolated as previously described^53^ with modifications. Briefly, 10 cm segments of small intestine were excised, opened longitudinally, and washed to remove luminal contents. Peyer’s patches and residual fat were removed, and tissue was cut into ∼5 mm pieces. Samples were sequentially washed in HBSS (Sigma Aldrich / H9394) and incubated in HBSS containing 2 mM EDTA (VWR / E177) to remove epithelial fractions, followed by multiple agitation cycles until supernatants became clear. Tissue was then digested in RPMI containing 20% FBS (gibco / 17479633), collagenase A (1mg/mL), and DNase I (0.05 mg/mL) for 30 min at 37 °C with shaking. The resulting suspension was filtered through 100 µm and 40 µm strainers, washed, and centrifuged (600 × g, 6 min, 4 °C). Cells were resuspended in FACS buffer supplemented with DNase I to reduce aggregation and processed for counting and downstream flow cytometry analysis.

#### Skin single-cell preparation

Dorsal skin was harvested from shaved and depilated 4 × 4 cm dorsum sections. Subcutaneous fat was removed after brief incubation in cold RPMI, and tissue was weighed prior to processing. Samples were minced into ∼1 mm² pieces and digested in 4 mL digestion buffer (RPMI containing collagenase A (1.7 mg/mL) and DNase I (100 µg/mL) for 1.5 h at 37 °C with intermittent vortexing every 30 min. Cell suspensions were mechanically dissociated using an 18G needle, followed by sequential filtration through 100 µm and 40 µm strainers with FACS buffer washes to remove debris and hair. Cells were centrifuged (700 × g, 8 min), washed, and resuspended in FACS buffer. The resulting single-cell suspension was used for downstream flow cytometry.

#### Testis single-cell preparation

Testis were extracted and mechanically minced with scissors. Samples were digested in 0.5 mL digestion buffer (collagenase A (1.5 mg/mL) and DNase I (0.1 mg/mL) for 20 min at 37 °C with shaking. Following enzymatic digestion, suspensions were mechanically dissociated and then filtered through a 70 µm strainer into 50 mL tubes and washed with FACS buffer. Cells were centrifuged (350 × g, 10 min, 4 °C), washed and resuspended in FACS buffer. The resulting single-cell suspension was used for counting and downstream flow cytometric analysis.

#### Bone marrow single-cell preparation

Hind limb bones were dissected from mice, cleaned and maintained in PBS on ice. Bone marrow was isolated by centrifugation using perforated 0.5 mL microcentrifuge tubes placed within 1.5 mL collection tubes. Following centrifugation (10,000 × g, 15 s) the collected bone marrow was resuspended and filtered through a 70 µm cell strainer into a 50mL falcon. The cell suspension was RBC lysed followed by centrifugation (500 × g, 5 min). Finally, cells were washed and resuspended in complete RPMI medium and used for counting and downstream applications.

### Flow cytometric staining

In short single cell suspensions were washed in PBS and stained with viability stain for 45 min at 4 °C. Following washing with FACS buffer, cells were incubated with anti-CD16/32 and True-Stain Monocyte Blocker™ (BioLegend / 426102) for 30 min at 4 °C. Surface staining was performed using antibody master mix for 45 min at 4 °C, followed by washing in FACS buffer. Where required, cells were subsequently incubated with secondary antibody mix for 15 min at 4 °C and washed again. If intracellular staining was needed, cells were fixed and permeabilized in Permeabilization/Fixation according to cenor recommendation, otherwise fixed with 1% formaldehyde (WVR / 11699455) for 20min. Stain mixes containing more than one than one Brillian violet dyes where mixed with 10% BD Horizon™ Brilliant Stain Buffer Plus (BD / 566385). The stained samples were acquired on the Cytek Aurora (3 laser 16V-14B-8R), and the fcs files were analysed using Flowjo v10. The antibodies used are listed in Suppl. Tables 1-4.

### BMDC culture and stimulation

Bone marrow cells were cultured in RPMI medium supplemented with 10% FBS, 0.2% Primocin (Invivogen / ant-pm-05), and 4% Flt3L-supernatant from own production with transfected CHO cell line. Cells were seeded at 10^6^ cells/ml in 12-well plate and incubated at 37 °C with 5% CO₂. On days 3 and 6, cultures were fed by replacing 0.2 mL of medium with fresh medium containing Flt3L.

On day 8, BMDCs were harvested, counted, and replated in flat-bottom 96-well plates at 2 × 10^5^ cells in 200 µL per well. Cells were stimulated as indicated in table 5 for 18 h before downstream analysis.

### OT-I/OT-II T-cell isolation and co-culture

Spleens and lymph nodes (inguinal, cervical, and axillary) were isolated from OT-I or OT-II mice and mechanically dissociated in cold PBS. Cell suspensions were filtered through 70 µm strainers and centrifuged (500 × g, 5 min, 4 °C). Splenocytes were subjected to RBC lysis for 5 min on ice, washed in FACS buffer, and combined with lymph node cells prior to counting. CD4^+^ or CD8^+^ T cells were isolated by magnetic positive selection using biotinylated antibodies and streptavidin nanobeads according to the manufacturer’s protocol. Purified cells were washed, resuspended in FACS buffer, and counted. Where indicated, T cells were labeled with proliferation dye (Tag-it Violet™ Proliferation and Cell Tracking Dye; Biolegend / 425101) for 20 min at room temperature, quenched with complete RPMI medium, washed, and resuspended for downstream applications.

For co-culture assays, 5 × 10^4^ T cells were cultured with 1 × 10^4^ BMDCs previously pulsed with OVA (200 µg/ml; Sigma-Aldrich, A5503-1G) for 18 h, as described above, in complete RPMI medium in 96-well U-bottom plates and incubated at 37 °C with 5% CO₂ for up to 4 days. For flow cytometric analysis, cells were harvested on day 2 and centrifuged (500 × g for 7 min at 4 °C). Cells were stained with an antibody cocktail (Table 4) for 20 min at 4 °C, washed with FACS buffer, and analyzed by flow cytometry.

**Table 1:** Tissue stain.

| **Antibody** | **Fluorochrome** | **Clone** | **Vendor** | **Concentration** | **Staining location** |
| --- | --- | --- | --- | --- | --- |
| CD103 | PE/Fire 810 | QA17A24 | BioLegend | 0,6667 µg/ml | Surface |
| CD11b | APC/Fire 810 | M1/70 | BioLegend | 0,0625 µg/ml | Surface |
| CD11c | BV650 | N418 | BioLegend | 0,125 µg/ml | Surface |
| CD135 (FLT3) | Biotin | A2F10 | BioLegend | 5 µg/ml | Surface |
| CD14 | BV785 | Sa14-2 | BioLegend | 0,0625 µg/ml | Surface |
| CD16/32 | BV711 | 93 | BioLegend | 1:400 | Surface |
| CD172a (SIRPα) | APC | QA20A06 | BioLegend | 1 µg/ml | Surface |
| CD19 | PE/Cyanine5 | 6D5 | BioLegend | 0,03125 µg/ml | Surface |
| CD192 (CCR2) | PE/Fire 640 | SA203G11 | BioLegend | 2 µg/ml | Surface |
| CD209a | BV480 | 5H10 | BD | 2 µg/ml | Intracellular |
| CD24 | PerCP-eFluor 710 | M1/69 | Invitrogen | 0,0625 µg/ml | Surface |
| CD26 (DPP-4) | FITC | H194-112 | BioLegend | 1,25 µg/ml | Surface |
| CD3 | PE/Cyanine5 | 17A2 | BioLegend | 0,0625 µg/ml | Surface |
| CD301b (MGL2) | PerCP/Cyanine5,5 | URA-1 | BioLegend | 2 µg/ml | Surface |
| CD45 | Alexa Fluor 532 | 30-F11 | Invitrogen | 0,5 µg/ml | Surface |
| CD45R/B220 | BV750 | RA3-6B2 | BioLegend | 0,5 µg/ml | Surface |
| CD64 (FcγRI) | BV421 | W18349C | BioLegend | 1 µg/ml | Surface |
| CD86 | Alexa Fluor 647 | GL-1 | BioLegend | 0,15625 µg/ml | Surface |
| CD88 (C5aR) | PE/Cyanine7 | 20/70 | BioLegend | 0,25 µg/ml | Surface |
| ESAM | PE | 1G8/ESAM | BioLegend | 0,25 µg/ml | Surface |
| F4/80 | APC/Fire 750 | BM8 | BioLegend | 1 µg/ml | Surface |
| Ly-6C | BV570 | HK1,4 | BioLegend | 0,25 µg/ml | Surface |
| Ly-6G | Alexa Fluor 700 | 1A8 | BioLegend | 0,625 µg/ml | Surface |
| MHC II (I-A/I-E) | Pacific Blue | M5/114,15,2 | BioLegend | 0,3125 µg/ml | Surface |
| NK1,1 | PE/Cyanine5 | PK136 | BioLegend | 0,125 µg/ml | Surface |
| Siglec-H | RB744 | 440c | BD | 1 µg/ml | Surface |
| Streptavidin | BV605 |  | BioLegend | 0,5 µg/ml | Surface |
| TER-119 | PE/Cyanine5 | TER-119 | BioLegend | 0,25 µg/ml | Surface |
| XCR1 | PE/Dazzle 594 | ZET | BioLegend | 0,0625 µg/ml | Surface |
| Zombie NIR<br>Fixable Viability Kit | Zombie NIR |  | BioLegend | 1:2000 | Surface |

**Table 2:** BM stain.

| **Antibody** | **Fluorochrome** | **Clone** | **Vendor** | **Concentration** | **Staining location** |
| --- | --- | --- | --- | --- | --- |
| CD115 (CSF-1R) | BV605 | AFS98 | BioLegend | 2 µg/ml | Intracellular |
| CD117 (c-kit) | BV785 | ACK2 | BioLegend | 0,25 µg/ml | Surface |
| CD11b | Alexa Fluor 660 | M1/70 | Invitrogen | 0,0625 µg/ml | Surface |
| CD11c | BV570 | N418 | BioLegend | 1:100 | Surface |
| CD127 (IL-7Rα) | APC/Fire 810 | S18006K | BioLegend | 0,5 µg/ml | Surface |
| CD135 (FLT3) | Biotin | A2F10 | BioLegend | 5 µg/ml | Surface |
| CD150 (SLAMF) | PerCP-eFluor 710 | mShad150 | Invitrogen | 0,5 µg/ml | Surface |
| CD16/32 | BV711 | 93 | BioLegend | 0 µg/ml | Surface |
| CD16/32 | BV711 | 93 | BioLegend |  | Surface |
| CD170 (Siglec-F) | APC | S17007L | BioLegend | 0,25 µg/ml | Surface |
| CD172a (SIRPα) | PerCP/Cyanine5,5 | QA20A06 | BioLegend | 2 µg/ml | Surface |
| CD19 | PE/Cyanine5 | 6D5 | BioLegend | 0,03125 µg/ml | Surface |
| CD192 (CCR2) | PE/Fire 640 | SA203G11 | BioLegend | 2 µg/ml | Surface |
| CD24 | Alexa Fluor 700 | M1/69 | BioLegend | 0,625 µg/ml | Surface |
| CD26 (DPP-4) | PE/Cyanine7 | H194-112 | BioLegend | 0,66667 µg/ml | Surface |
| CD3 | PE/Cyanine5 | 17A2 | BioLegend | 0,0625 µg/ml | Surface |
| CD34 | BV421 | SA376A4 | BioLegend | 4 µg/ml | Surface |
| CD45 | Alexa Fluor 532 | 30-F11 | Invitrogen | 0,25 µg/ml | Surface |
| CD45R/B220 | BV750 | RA3-6B2 | BioLegend | 0,25 µg/ml | Surface |
| CD48 | PE/Dazzle 594 | HM48-1 | BioLegend | 0,083333 µg/ml | Surface |
| CD86 | Alexa Fluor 647 | GL-1 | BioLegend | 0,15625 µg/ml | Surface |
| CD88 (C5aR) | BUV737 | 20/70 | BD | 2 µg/ml | Surface |
| F4/80 | PE/Cyanine5 | BM8 | BioLegend | 0,5 µg/ml | Surface |
| FcεR1α | PE | MAR-1 | BioLegend | 0,125 µg/ml | Surface |
| Ly-6A/E (Sca-1) | BV650 | D7 | BioLegend | 0,25 µg/ml | Surface |
| Ly-6C | PE/Fire 810 | HK1,4 | BioLegend | 0,0625 µg/ml | Surface |
| Ly-6D | FITC | 49-H4 | BioLegend | 0,078125 µg/ml | Surface |
| Ly-6G | BV480 | 1A8 | BD | 0,5 µg/ml | Surface |
| MHC II (I-A/I-E) | Pacific Blue | M5/114,15,2 | BioLegend | 0,3125 µg/ml | Surface |
| NK1,1 | PE/Cyanine5 | PK136 | BioLegend | 0,125 µg/ml | Surface |
| Siglec-H | RB744 | 440c | BD | 1 µg/ml | Surface |
| Streptavidin | APC/Fire 750 |  | BioLegend | 0,5 µg/ml | Surface |
| TER-119 | PE/Cyanine5 | TER-119 | BioLegend | 0,25 µg/ml | Surface |
| Zombie NIR<br>Fixable Viability Kit | Zombie NIR |  | BioLegend | 1:2000 | Surface |

**Table 3:** BMDC stain.

| **Antibody** | **Fluorochrome** | **Clone** | **Vendor** | **Concentration** | **Staining location** |
| --- | --- | --- | --- | --- | --- |
| CD103 | PE/Fire 810 | QA17A24 | BioLegend | 0,25 µg/ml | Surface |
| CD11b | APC/Fire 810 | M1/70 | BioLegend | 0,03125 µg/ml | Surface |
| CD11c | BV570 | N418 | BioLegend | 0,180556 | Surface |
| CD14 | BV650 | rmC5-3 | BD | 0,25 µg/ml | Surface |
| CD16/32 | BV711 | 93 | BioLegend | 1:400 | Surface |
| CD172a (SIRPα) | APC/Cyanine7 | P84 | BioLegend | 0,0625 µg/ml | Surface |
| CD197 (CCR7) | Alexa Fluor 700 | 4B12 | Invitrogen | 0,25 µg/ml | Surface |
| CD24 | PerCP-eFluor 710 | M1/69 | Invitrogen | 0,03125 µg/ml | Surface |
| CD26 (DPP-4) | APC | H194-112 | BioLegend | 0,5 µg/ml | Surface |
| CD273 (PD-L2) | RB744 | MIH37 | BD | 0,5 µg/ml | Surface |
| CD274 (PD-L1) | Alexa Fluor 660 | MIH5 | Invitrogen | 0,125 µg/ml | Surface |
| CD301b (MGL2) | PE/Dazzle 594 | URA-1 | BioLegend | 0,25 µg/ml | Surface |
| CD317 (BST2, PDCA-1) | Alexa Fluor 647 | 129C1 | BioLegend | 1,25 µg/ml | Surface |
| CD40 | Pacific Blue | 01,03,2023 | BioLegend | 1,25 µg/ml | Surface |
| CD45R/B220 | BV750 | RA3-6B2 | BioLegend | 0,0625 µg/ml | Surface |
| CD64 (FcγRI) | PE | S18017D | BioLegend | 0,0625 µg/ml | Surface |
| CD86 | BV605 | PO3 | BioLegend | 1 µg/ml | Surface |
| Ly-6C | RB545 | AL-21 | BD | 0,25 µg/ml | Surface |
| MHC II (I-A/I-E) | Spark Violet 538 | M5/114,15,2 | BioLegend | 1,25 µg/ml | Surface |
| Streptavidin | FITC |  | BioLegend | 1,25 µg/ml | Surface |
| XCR1 | Biotin | ZET | BioLegend | 1,25 µg/ml | Surface |
| Zombie NIR Fixable Viability Kit | Zombie NIR |  | BioLegend | 1:2000 | Surface |

**Table 4:** OT staining.

| **Antibody** | **Fluorochrome** | **Clone** | **Vendor** | **Concentration** | **Staining location** |
| --- | --- | --- | --- | --- | --- |
| 7-AAD Viability Staining Solution | 7-AAD |  | BioLegend | 0,5 µg/ml | Surface |
| CD3ε | BV785 | 145-2C11 | BioLegend | 2 µg/ml | Surface |
| CD4 | BV650 | RM4-5 | BioLegend | 0,5 µg/ml | Surface |
| CD8a | APC | S18018E | BioLegend | 0,5 µg/ml | Surface |

**Table 5:**
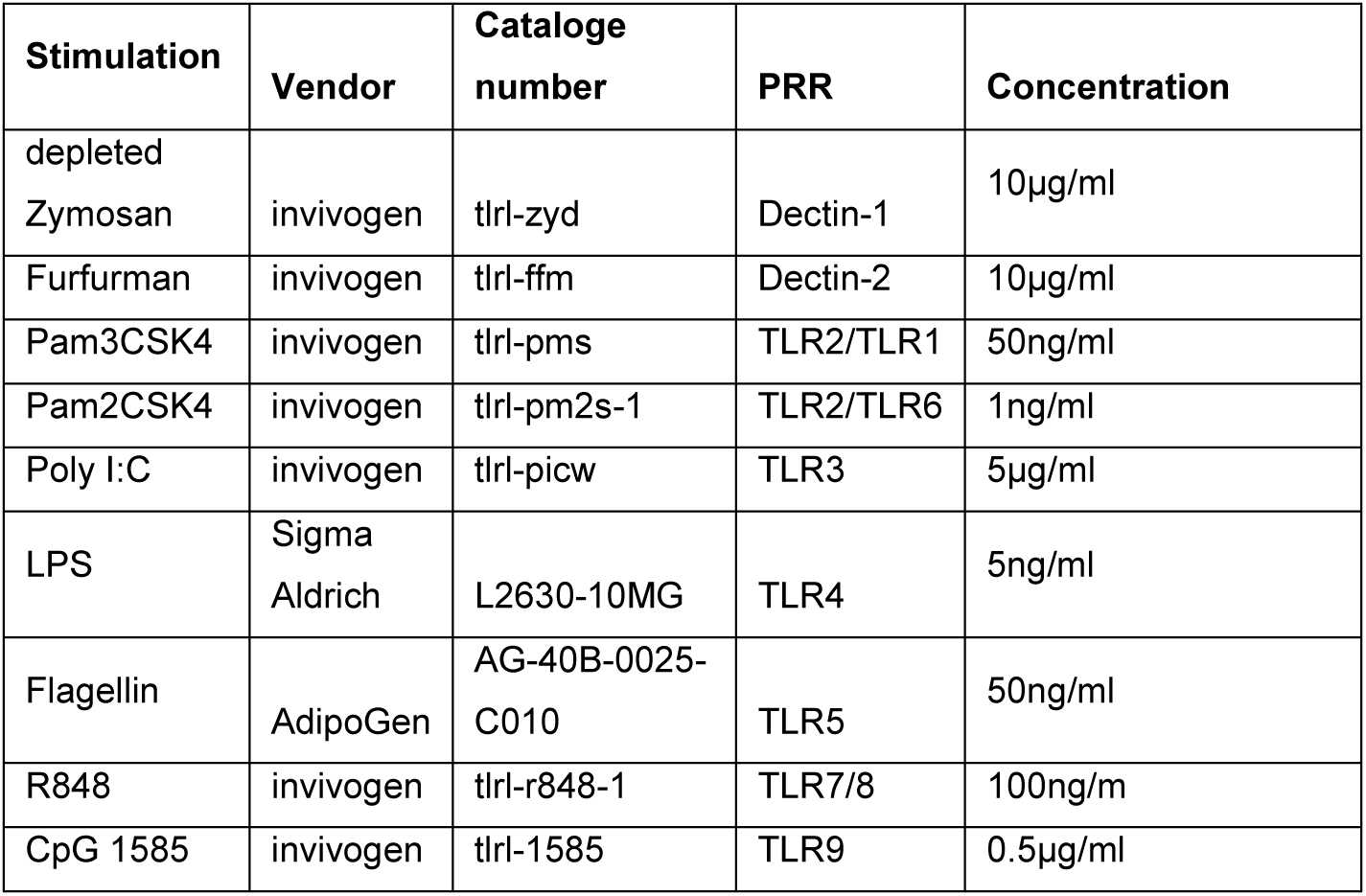
Stimualtions for BMDC culture.

### Single-cell RNA-seq data processing and analysis

#### Dataset curation and reanalysis

For single-cell RNA sequencing analysis, cells from the intestine, lung, spleen, and liver in dataset GSE153562 were reanalysed. All downstream analyses were performed in the R environment. During quality control, cells with a mitochondrial gene percentage greater than 15% were excluded. Ambient RNA contamination was estimated using the DecontX package, and cells with an ambient RNA contamination score greater than 0.5 were removed. In addition, doublet detection was performed for each sample using DoubletFinder, and predicted doublets were excluded from further analysis. Data preprocessing was conducted using the SCTransform workflow implemented in the Seurat package. Samples were subsequently integrated using Harmony for batch correction and downstream joint analysis. Cell clusters were annotated based on established marker genes listed in table 6. Only dendritic cell populations, including cDC1, cDC2, pDC, and migratory dendritic cells, were retained for subsequent analyses. For dimensionality reduction and visualization, Uniform Manifold Approximation and Projection was applied based on the Harmony-corrected embedding. To quantitatively evaluate age-associated dynamics of senescence-related markers, pseudobulk expression profiles were generated by aggregating expression values at the sample level within each relevant cell population. The proportion of Cdkn1a-expressing cells was calculated as the percentage of cells with detectable Cdkn1a expression, defined as expression values greater than zero.

**Table 6.**
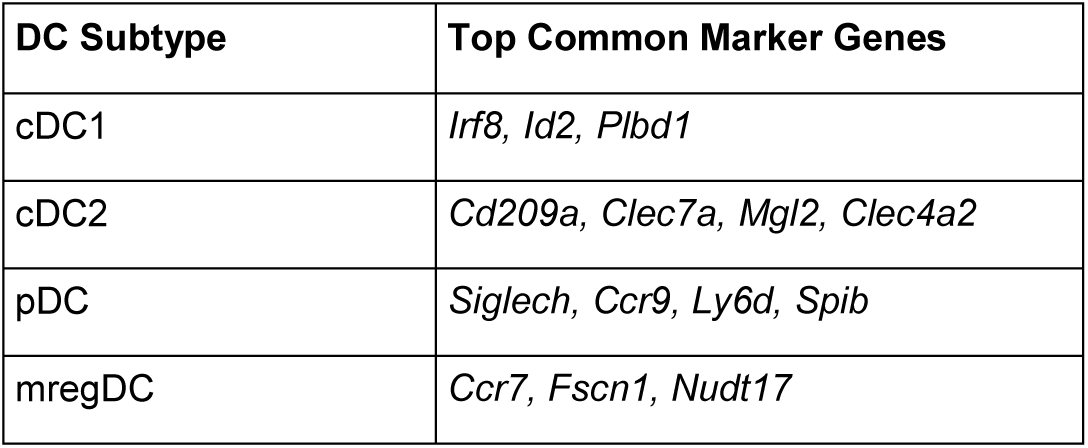
Top differentially expressed marker genes across dendritic cell subsets in lung, liver, small intestine tissues (GSE153562)

#### Senescence-associated transcriptomic shift analysis (p21-shift)

To identify transcriptional changes associated with Cdkn1a expression in immune cell populations, a targeted differential expression analysis, referred to as p21-shift analysis, was performed. Within each tissue and dendritic cell subtype, cells were classified according to endogenous Cdkn1a expression. Cells with detectable Cdkn1a expression were defined as p21-positive, whereas cells without detectable Cdkn1a transcripts were defined as p21-negative. Comparisons with fewer than three cells in either group were excluded from further analysis. Differential expression between p21-positive and p21-negative cells was assessed using the Wilcoxon rank-sum test implemented in the FindMarkers function of Seurat. To evaluate the conservation and heterogeneity of Cdkn1a-associated transcriptional signatures across tissues and dendritic cell subsets, DEG lists from different tissues and cell types were intersected using the UpSetR package. The top 30 intersections were visualized to identify shared pan-tissue signatures as well as tissue- or subtype-specific gene responses.

#### Age-dependent alterations within the DC compartment

To assess whether the transcriptomic profile of dendritic cells changes with age, a secondary differential expression analysis was performed. For dataset GSE153562, cells from 6–8-week-old mice were classified as young, whereas cells from 18-month-old mice were classified as old. Intermediate or unmatched age groups, such as 12-month-old samples, were excluded from this comparison. Differential expression analysis comparing old versus young each organ and dendritic cell subtype using the Wilcoxon rank-sum test implemented in the FindMarkers function of Seurat. Genes with a Benjamini-Hochberg-adjusted p value < 0.05 and an absolute log2 fold change > 0.25 were considered significantly differentially expressed. Analyses were restricted to organ and dendritic cell subtype combinations containing at least three cells per age group; tissues not meeting this criterion were excluded from downstream analysis due to insufficient cell numbers.

### Statistical Analysis

Graphpad prism (version 8.5) was used for unpaired t test comparing MFI expression or percentages in defined cell types in flow cytometry data and R 4.5.1 for bioinformatics.

